# A new method for quantifying APT and NOE(-3.5) using chemical exchange saturation transfer with double saturation powers (DSP-CEST)

**DOI:** 10.1101/2022.11.13.516305

**Authors:** Yu Zhao, Casey Sun, Zhongliang Zu

**Affiliations:** Vanderbilt University Institute of Imaging Science, Nashville, US; Department of Radiology and Radiological Sciences, Vanderbilt University Medical Center, Nashville, US; Department of Radiology, West China Hospital of Sichuan University, Chengdu, China; Department of Chemistry, University of Florida, Gainesville, US

**Keywords:** chemical exchange saturation transfer (CEST), amide proton transfer (APT), nuclear Overhauser enhancement, exchange rate, tumor

## Abstract

**Purpose:** Quantifications of amide proton transfer (APT) and nuclear Overhauser enhancement (NOE(−3.5)) mediated transfer with high specificity are challenging since their signals measured in a Z-spectrum are overlapped with confounding signals from direct water saturation (DS), semi-solid magnetization transfer (MT) and chemical exchange saturation transfer (CEST) of fast-exchange pools. In this study, based on two canonical CEST acquisitions with double saturation powers (DSP), a new data-postprocessing method is proposed to specifically quantify the effects of APT and NOE.

**Methods:** For CEST imaging with relatively low saturation powers 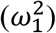, both the fast-exchange CEST effect and the semi-solid MT effect increase linearly with 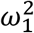 whereas the slow-exchange APT/NOE(−3.5) effect has no such a dependence on 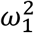, which is exploited to isolate the APT and NOE effects from the confounding signals in this study. After a mathematical derivation for the establishment of the proposed method, numerical simulations based on Bloch equations are then performed to demonstrate its specificity to detections of the APT and NOE effects. Finally, an *in vivo* validation of the proposed method is conducted using an animal tumor model at a 4.7-T MRI scanner.

**Results:** The simulations show that DSP-CEST can quantify the effects of APT and NOE and substantially eliminate the confounding signals. The in vivo experiments demonstrate that the prosed DSP-CEST method is feasible for the imaging of tumors.

**Conclusion:** The data-postprocessing method proposed in this study can quantify the APT and NOE effects with considerably increased specificities and a reduced cost of imaging time.

## INTRODUCTION

Chemical exchange saturation transfer (CEST) MRI leverages a sensitivity enhancement mechanism to indirectly detect dilute mobile molecules that contain exchangeable/coupling protons (1–5). In CEST imaging, the exchangeable/coupling protons are saturated by frequency-selective saturation RF pulses and subsequently transferred to the bulk water via chemical exchange or dipolar interaction, yielding a reduced signal of the bulk water. Usually, the frequency offset of saturation RF pulse is varied to acquire a Z-spectrum that plots the labeled water signals as a function of the frequency offset such that multiple types of exchangeable/coupling protons that are associated with different molecules or chemical groups and thus possess distinct chemical shifts can be detected. Amide proton transfer (APT), a variant of CEST imaging, seeks to detect the backbone amide of mobile proteins/peptides as CEST signals at 3.5 ppm (6), which has demonstrated great potential in diagnosing tumors (7–10), ischemic stroke (11–13), multiple sclerosis (14,15), traumatic brain injury (16), Alzheimer’s disease (17–21) and Parkinson’s disease (22,23), etc. The APT imaging relies on slow chemical exchanges and thus can be conducted with low saturation RF powers (e.g., B_1_ < 1 μT). Another variation of CEST imaging that also relies on the slow exchange/coupling is Nuclear Overhauser enhancement (NOE) mediated saturation transfer MRI, which measures the CEST signals at approximately −3.5 ppm from mobile macromolecules with finite linewidth (e.g., large proteins, phospholipids) (24–28). In previous studies, the NOE-mediated saturation transfer effect has been applied to brain tumors (28–31), Alzheimer’s disease (32) and spinal cord injury (33), etc.

To date, detections of the APT effect and the NOE effect with high specificity are still challenging since their CEST signals are overlapped with background signals from the direct water saturation (DS), the semi-solid magnetization transfer (MT) and the fast-exchange saturation. Although the DS signals are weak at low saturation powers, the MT-sourced signals are still considerable. In addition, the amine on glutamate and protein-contained lysine with a chemical shift of 3 ppm (34,35) could contribute substantial CEST signals at 3.5 ppm and thus confound the APT signals because the fast exchange between amine and water leads to a broad line-shape of the amine CEST signals. Notably, our previous study suggests that the fast-exchange amine has contributions to the CEST signals at 3.5 ppm even at relatively low saturation powers (36).

To isolate the APT and NOE effects from the confounding signals, a reference signal that has contributions from the background sources and is relatively insensitive to the APT or NOE effects is usually first obtained and then subtracted with the label signal. However, the acquisition of an accurate reference in a complex biological system is challenging. For example, an asymmetric analysis is used to quantify the amide exchange in earlier studies, which is found to be suffering from a reference with the contamination from NOE on the opposite side relative to the water peak in the Z-spectrum (37). Multiple-pool Lorentzian fit, that attempt to separate the effects sourced from the amide, amine, water, NOEs, and semi-solid MT pools, has also been used to quantify the APT and NOE effects (28,38–40). However, its accuracy strongly depends on the image SNR, initial values, boundary conditions, and the number of fitting pools as well as their line shapes. At lower fields, it would be more difficult to separate these CEST peaks since some of them could more seriously overlap with each other. Besides, Lorentzian difference (LD) analysis (29) and extrapolated semi-solid MT reference (EMR) (41), that seek to obtain the reference signal by fitting the DS and MT effects, has also been developed to quantify the CEST and NOE effects. Note that, although the two methods cannot separate the amine-exchange contamination from the APT signals, they have advantages over the fitting methods at relatively low fields (with reduced frequency separations), considering that the overlapping of CEST peaks is not needed to be coped with. In addition to these data-postprocessing methods, other methods based on complex exchange-editing RF pulses and readout sequences, such as variable delay multi-pulse (VDMP) (42–44), frequency labeled exchange transfer (FLEX)) (45), chemical exchange rotation transfer (CERT) (37,46–48), frequency alternating RF irradiation (SAFARI) (49,50) and length and offset varied saturation (LOVARS) (51,52), have also been demonstrated to be able to separately quantify CEST effects with different signal sources, whereas they have limited applications due to sophisticated designs of pulse sequences.

In this study, we propose a new data-postprocessing method to specifically quantify the APT and NOE effects based on two canonical CEST-MRI acquisitions with double saturation powers (DSP), where the signal acquired at a low saturation power is directly used for the label signal and the signal acquired at a high saturation power is processed by a well-tailored mathematical transformation to create a reference signal. Firstly, a mathematical modeling with approximations is used to interpret the proposed method with physical pictures. Then, numerical simulations based on Bloch equations without the approximations are performed to demonstrate its specificity to detections of the APT and NOE effects. Finally, an *in vivo* validation of the proposed method is conducted using an animal tumor model at a 4.7-T MRI scanner.

## THEORY

For CEST imaging with continuous RF radiation at a low power, the CEST signal (*S*) measured in a steady state can be modeled in a concise formation with a comprehensive relaxation rate (*R*_l*ρ*_(*ω_RF_*)) that is defined in a rotating frame and dependent on the frequency offset (*ω_RF_*) of the saturation RF relative to the water resonance frequency, and it is described as

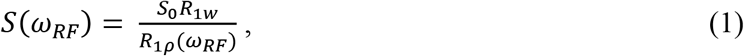

where *S*_0_ is the control signal measured without the RF saturation; *R*_1*w*_ is the longitudinal relaxation rate of the water. In this formation, the comprehensive relaxation includes CEST/NOE effects from the slow and fast exchange/coupling (e.g., the slow-exchange pools: APT, NOE; the fast exchange pool: amine CEST)) along with the DS and MT effects, which can be expressed as the summation of separable relaxations of the corresponding pools if their relative concentrations are much lower than 1 (53,54),

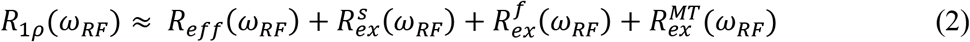

where 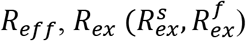, and 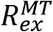 correspond to the water longitudinal relaxation, the CEST effect from the slow exchange/coupling pool (the ‘s’ superscript) and the fast exchange pool (the ‘f’ superscript), and the semi-solid MT effect in the rotating frame, respectively. Based on previous studies (53,54), 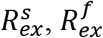 and *R_eff_* can be further derived as

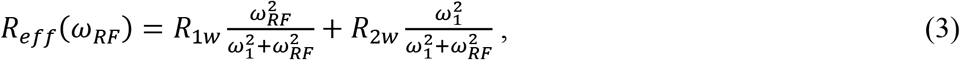

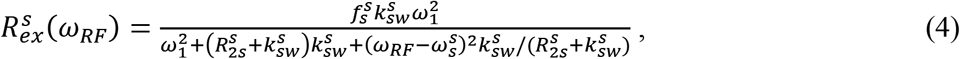

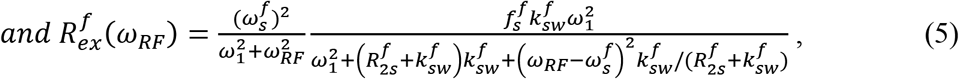

where *f_s_* is the relative concentration of the solute; *R*_1_ and *R*_2_ are the longitudinal and transverse relaxation rate, respectively; *ω*_1_ denotes the saturation RF power; *k_sw_* is the solute-to-water exchange rate; *ω_s_* is the frequency offset of the solute relative to the water resonance frequency; the ‘w’ and ‘s’ subscripts denote the water and the solute, respectively; and the ‘s’ and ‘f’ superscripts denote the slow exchange/coupling pool and the fast exchange pool, respectively. Note that *ω*_1_ = *γB*_1_, where *γ* is the gyromagnetic ratio of proton and *B*_1_ is the amplitude of the RF field.

Then, Eq. (3) and Eq. (5) can be approximated to obtain their concise formations under the condition of low saturation powers. Specifically, when the frequency offset of the APT/ NOE(−3.5) (i.e., 2815rad/s and 4398rad/s at 3 T and 4.7 T, respectively) is much larger than *ω*_1_ (e.g., *ω*_1_ = 268rad/s for B_1_ = 1 μT), R_eff_ can be approximated as

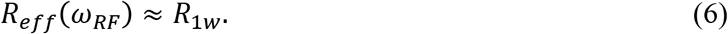

For fast exchange pools, *k_sw_* (e.g., *k_sw_* = 5000 s^−1^ for the amine (55)) is also much larger than the saturation power and thus *ω*_1_H_ in the denominator of Eq. (5) can be ignored and Eq. (5) can be then approximated as

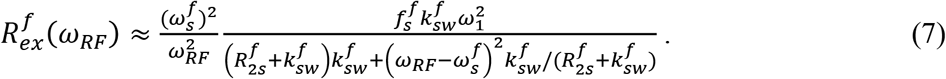

In this study, based on two measurements under different low saturation powers (*B*_1_ less than 1 μT), new metrics are constructed to isolate the slow exchange/coupling effects at ±3.5 ppm as below. Based on Eq. (1), Eq. (2) and Eq. (6), the CEST signal (S_L_) measured with a relatively low saturation power (*ω*_1_L_) can be approximately modeled as

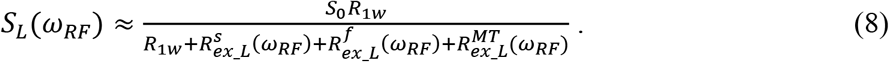

Then, we construct an reference signal based on the CEST signal (*S_H_*) measured with a relatively high saturation power (*ω*_1_H_), the control signal (*S*_0_) and a prior parameter (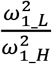, determined in MRI experiments), which is defined as

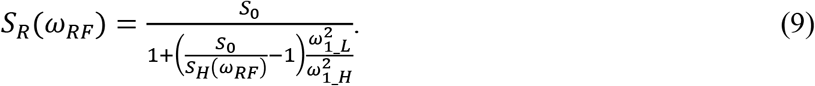

Note that *S_R_*(*ω_RF_*) is a single -variable function of the measured signal *S_H_*. By multiplying both the numerator and the denominator of Eq. (9) by *R*_1*w*_, then substituting *S_H_*(*ω_RF_*) with 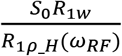 (according to Eq. (1)), and subsequently expanding *R*_1*ρ_H*_ according to Eq. (2), Eq. (9) can be transformed as

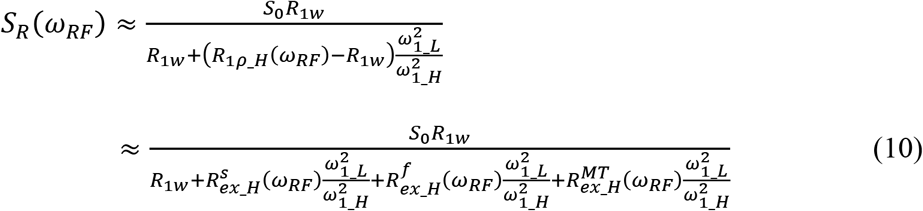

According to Eq. (7) and APPENDIX A, 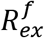 and 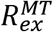 are approximately proportional to 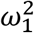, and thus we have 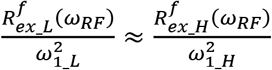 and 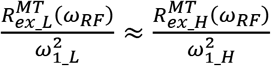. Hence, Eq. (9) can be further transformed as

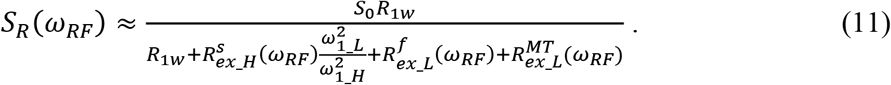

By comparing Eq. (8) with Eq. (11), it can be inferred that the difference between *S_R_* and *S_L_* measured in the DSP-CEST can be used to measure the slow-exchange effects with the insusceptibility to the fast-exchange and MT effects. Magnetization transfer ratio (MTR) and apparent exchange-dependent relaxation (AREX) (53,54,56) are two widely used metrics to process the label and reference signals to quantify CEST effect. Here, we also apply the two metrics to the label and reference signals to quantify the APT/NOE effect,

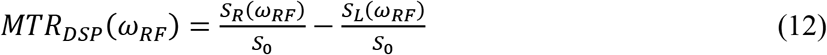

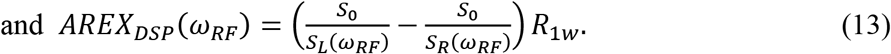

Note that the specificity of the two metrics to detections of the slow-exchange effect will be further validated using numerical simulations (see the next section), which do not rely on the approximations mentioned above.

## MATERIALS AND METHODS

### Numerical simulations

Based on the coupled Bloch equations, numerical simulations for a five-pool model (amide + amine + water + NOE(−3.5) + semi-solid MT) were performed to validate the proposed DSP-CEST approach. Numerical calculations of Bloch equations were conducted using the ordinary differential equation solver (ODE45) in MATLAB (Math works, Natick, MA, USA), which is well described in a previous study (36). The parameters of the continuous saturation RF are a duration of 5 seconds and two saturation powers of B_1_*L*_ = 0.5 μT and B_1_*H*_ = 1 μT. In the numerical simulations, the relevant parameters were varied to investigate their influences on the DSP-CEST signal in the simulations as below.

1. To evaluate the capability of the DSP-CEST to separate the slow-exchange CEST effect (i.e., APT and NOE) from the fast-exchange amine CEST effect and the semi-solid MT effect, simulations were performed with varied concentrations of the semi-solid pool (*f_m_*= 5%, 10% and 15%) and varied concentrations of the fast-exchange amine (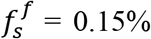, 0.3%, and 0.45%), individually. Note that other sample parameters for these simulations were kept constant as shown in Table 1. Then, MTR_DSP_ and AREX_DSP_ spectra were obtained using Eq. (12) and Eq. (13). In addition, the corresponding R_ex_ spectra for the fast-exchange amine were calculated using Eq. (5) with B_1_*L*_ = 0.5 μT and then compared with the AREX_DSP_ spectra.
2. To investigate the dependence of the DSP-CEST signals (defined by MTR_DSP_ and AREX_DSP_) on the concentration and the exchange/coupling rate of the target pools (APT and NOE), we performed the simulations with the following parameters: (1) for APT, 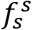 varied from 0.04 % to 0.2 % and *k_sw_* = 50 s^−1^; (2) for APT, *k_sw_* varied from 10 s^−1^ to 500 s^−1^ and 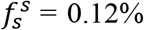; (3) for NOE, 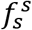 varied from 0.5% to 2.5% and *k_sw_* = 10 s^−1^; (4) for NOE, *k_sw_* varied from 1 s^−1^ to 50 s^−1^ and 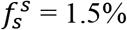. The four sets of sample parameters are combined with the varied concentration of the fast-exchange amine pool and semi-solid MT pool mentioned above, which are used to further investigate the robustness of the proposed method against the effects from the confounding pools. Also note that other sample parameters for these simulations were kept constant as shown in Table 1.
3. To investigate the influences of the *B*_0_ and *B*_1_ inhomogeneity on the DSP-CEST signals of the APT and the NOE, the simulations are performed with the *B*_0_ inhomogeneity spanning from −40 Hz to 40 Hz and the *B*_1_ inhomogeneity spanning from 80 % to 120 %, individually. Note that the *B*_1_ inhomogeneity is defined by the ratio of the deviated amplitude of RF field to the nominal value.

**Table 1.** Sample parameters used in the numerical simulations.

| Parameters | water | amide | fast exchange amine | NOE(-3.5) | Semi-solid MT |
| --- | --- | --- | --- | --- | --- |
| $f_s$ (%) | 100 | 0.12 | 0.3 | 1.5 | 10 |
| $k_{sw}$ ($s^{-1}$) | - | 50, 100 | 5000<sup>a</sup> | 10, 20 | 25 |
| $T_1$ (s) | 1.5 | 1.5 | 1.5 | 1.5 | 1.5 |
| $T_2$ (ms) | 60 | 2 | 10 | 0.5 | 0.015 |
| $\Delta$ (ppm) | 0 | 3.5 | 3 | -3.5 | 0 |

<sup>a</sup> 5000-7000 $s^{-1}$ in Ref (67,68)

### Animal Preparation

Three rats bearing 9L tumors were used for the *in vivo* validation of the proposed method in this study. Each rat was injected with 1 × 10^5^ 9L glioblastoma cells in the right brain hemisphere for the induction of brain tumor, and then imaged after 2 to 3 weeks. All rats were immobilized and anesthetized with 2-3% isoflurane and 97-98% oxygen during the experiments. Respiration rate was monitored to be in a range from 40 to 70 breaths per minute. Rectal temperature was maintained at 37°C using a warm-air feedback system (SA Instruments, Stony Brook, NY). All animal procedures were approved by the Animal Care and Usage Committee of Vanderbilt University Medical Center.

### MRI

In the DSP-CEST, after the signal preparation based on a rectangular RF saturation pulse with a duration of 5 seconds, single-shot spin-echo echo planar imaging (SE-EPI) is used for a 2D image readout with parameters: matrix size = 64 *×* 64, field of view = 30 *×* 30 mm^2^. DSP-CEST Z-spectra (B1 = 0.5 μT and 1 μT) were acquired with the frequency offsets from −2000 Hz to −1250 Hz with a step size of 250 Hz (−10 ppm to −6.25 ppm with a step size of 1.25 ppm at 4.7 T), −1000 Hz to 1000 Hz with a step size of 25 Hz (−5 ppm to 5 ppm with a step size of 0.125 ppm at 4.7 T), and 1250 Hz to 2000 Hz with a step size of 250 Hz (6.25 ppm to 10 ppm with a step size of 1.25 ppm at 4.7 T). Control images were acquired with the frequency offset of 100,000 Hz (500ppm at 4.7T). R_1w_ was obtained from an inversion recovery method. All measurements were performed on a Varian 4.7-T magnet with a 38-mm receive coil.

### Data analysis and statistics

In this study, the APT and NOE(−3.5) signals measured with the DSP-CEST were compared with those quantified with the LD analysis. In this analysis, the label signal is the measured signal and the reference signal (S_LD_R_) is estimated by a two-pool (water and semi-solid MT) model fitting of the CEST Z-spectrum with frequency offsets of ±2000, ±1750, ±1500, ±1250, ±100, ±75, ±50, ±25, and 0 Hz (−10 to −6.25ppm, −0.5 to 0.5ppm, and 6.25 to 10ppm at 4.7). Table 2 lists the starting points and boundaries of the LD analysis. The goodness of the LD fitting was assessed by the sum of squared errors. MTR_LD_ and AREX_LD_ spectra were obtained using Eq. (12) and Eq. (13) by replacing the S_DSP_R_ and S_DSP_L_ with S_LD_R_ and S_LD_L_. Finally, a statistical analysis of the CEST signals that are derived from the two methods was performed based on regions of interest (ROIs) that were delineated from the tumor region and the contralateral normal tissues in the rat brains bearing tumors. Student’s t-test was employed to evaluate the difference of the signals between tumors and contralateral normal tissues. It was considered to be statistically significant if *P* < 0.05.

**Table 2.** Starting points and boundaries of the amplitude (A), the peak width (W), and the chemical shift (Δ) of the water and semi-solid MT pools in the LD analysis. The unit of the peak width and the chemical shift t is ppm.

|  | Start | Lower | Upper |
| --- | --- | --- | --- |
| $A_{\text{water}}$ | 0.9 | 0.02 | 1 |
| $W_{\text{water}}$ | 1.4 | 0.1 | 10 |
| $\Delta_{\text{water}}$ | 0 | -1 | 1 |
| $A_{\text{semi-solid MT}}$ | 0.1 | 0 | 1 |
| $W_{\text{semi-solid MT}}$ | 25 | 10 | 100 |
| $\Delta_{\text{semi-solid MT}}$ | **0** | **-4** | **4** |

## RESULTS

In Fig. 1, numerical simulations demonstrate how the background signals from the MT and the fast-exchange amine are suppressed in the DSP-CEST imaging of the APT and NOE. In Fig. 1A and 1B, the conventional CEST Z-spectra with double saturation powers (B_1_L_ = 0.5 μT and B_1_H_ = 1 μT) and varied concentrations of the semi-solid MT pool (Fig. 1A) and the fast-exchange amine (Fig. 1B) are plotted with the raw data from the simulations. Fig. 1C and 1D show the corresponding DSP-CEST Z-spectra, where S_DPS_L_ is the raw data simulated with B_1_L_ = 0.5 μT and S_DPS_R_ is obtained from the mathematical transformation (in Eq. (9)) of the raw data simulated with B_1_L_ = 1 μT. In the DSP-CEST Z-spectra, the data points beyond the proximal frequencies of APT and NOE match well, suggesting that the difference of the DSP-CEST Z-spectra could be used to suppress the effects from the semi-solid MT and the fast-exchange amine. Fig. 1E-1H show the MTR_DPS_ and AREX_DPS_ spectra that are derived from the simulated DSP-CEST Z-spectra. Note that although the semi-solid MT effects change greatly with its concentration (see the CEST signals beyond ±5 ppm in Fig. 1A and 1C), the MTR_DPS_ at + 3.5ppm (Fig. 1F) and the AREX_DPS_ at ± 3.5 ppm (Fig. 1G and 1H) do not exhibit obvious dependencies on the changes in the MT effects. The small and non-ignorable changes of the MTR_DPS_ at −3.5ppm are observed in Fig. 1E, which should be due to the shine through effect from the semi-solid MT (36,53).

**Fig. 1.**
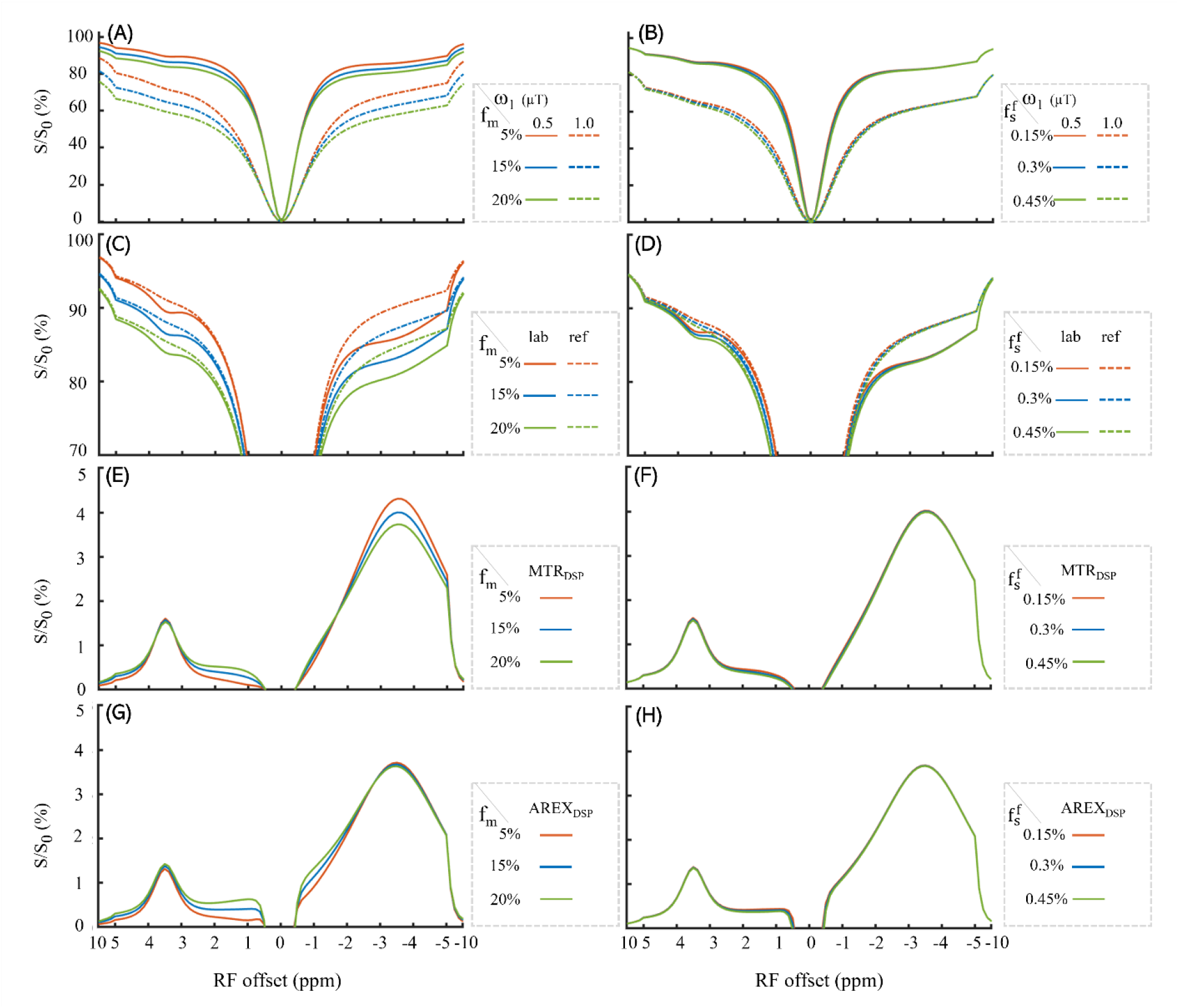
Simulated CEST Z-spectra with B_1_*L*_ = 0.5 μT and B_1_*H*_ = 1 μT (A, E), the corresponding S_DSP_L_ and S_DSP_R_ spectra (B, F), the MTR_DSP_ (C,G) and AREX_DSP_ (D, H) with three concentrations of the MT pool (left column) and three concentrations of the fast-exchange amine pool (right column). The numerical simulations is based on a five-pool model (amide + amine + water + NOE(−3.5) + semi-solid MT).

Fig. 2 and Fig. 3 show the influences of the MT and fast-exchange amine on the DSP-CEST signals of APT and NOE that are simulated with varied concentrations 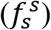 and varied exchange/coupling rates (*k_sw_*). The DSP-CEST signals of APT and NOE that are determined by MTR_DSP_ (Fig. 2) and AREX_DSP_ (Fig. 3) are plotted as a function of their concentration and exchange/coupling rate for different *f_m_* and 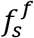. Note that the corresponding R_ex_ values (calculated using Eq. (5)) were also plotted for a comparison with the AREX_DSP_ in Fig. 3. Four important summaries can be made from Fig. 2 and Fig. 3: (1) both the APT and NOE(3.5) signals determined by MTR_DSP_ and AREX_DSP_ increase linearly with 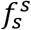, and first increased and then decreased with *k_sw_*. (2) the APT and NOE(3.5) signals determined by MTR_DSP_ are more susceptible to the influences of the MT and fast-exchange amine than those determined by AREX_DSP_; (3) the MT variations in a biophysically realistic conditions has more influences on the APT and NOE(3.5) signals determined by the two metrics than on the fast-exchange amine; (4) the decrease of the AREX_DSP_ values with increasing the exchange rate is greater than that of the R_ex_, and thus the DSP-CEST has an exchange-rate low-pass filter effect.

**Fig. 2.**
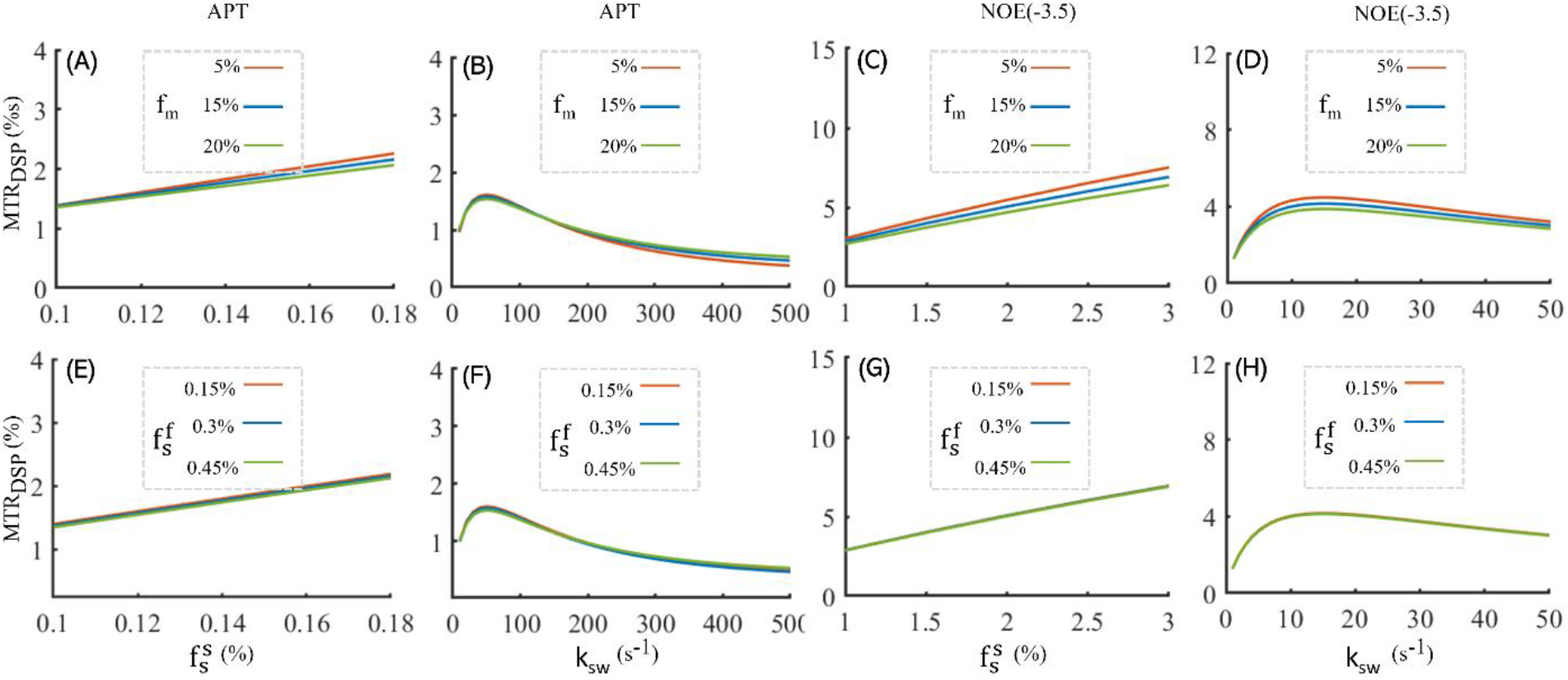
MTR_DSP_ signals of APT and NOE(−3.5) with varied concentrations 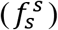 and varied exchange/coupling rates (*k_sw_*) simulated with three concentrations of the MT pool (1^st^ row) and three concentrations of the fast-exchange amine pool (2^nd^ row).

**Fig. 3.**
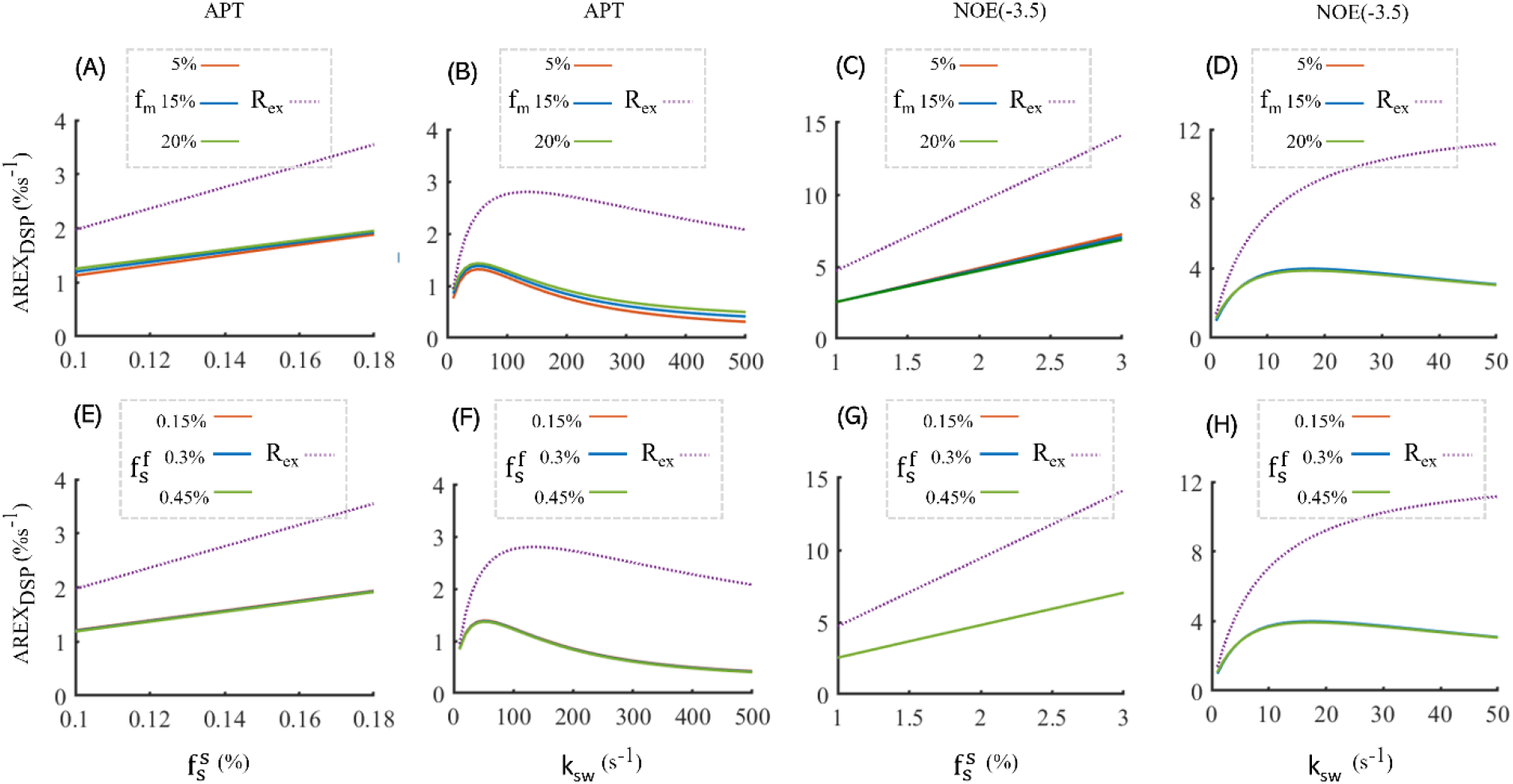
AREX_DSP_ signals of APT and NOE(−3.5) with varied concentrations 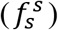 and varied exchange/coupling rates (*k_sw_*) simulated with three concentrations of the MT pool (1^st^ row) and three concentrations of the fast-exchange amine pool (2^nd^ row). Note that the corresponding R_ex_ (see the detailed definition in text) are also plotted.

In Fig. 4, the influences of the inhomogeneity of *B*_0_ and *B*_1_ on the MTR_DSP_ and AREX_DSP_ of APT and NOE(−3.5) are demonstrated using numerical simulations, where the ratio of the MTR_DSP_ or AREX_DSP_ measured with the inhomogeneities to the nominal MTR_DSP_ or nominal AREX_DSP_ are plotted as a function of the deviations of *B*_0_ and *B*_1_. The inhomogeneity of *B*_1_ (Fig. 4E-4H) has substantial influences on the MTR_DSP_ and AREX_DSP_ of APT and NOE while the inhomogeneity of *B*_0_ (Fig. 4A-4D) has less influences especially on the MTR_DSP_ and AREX_DSP_ of NOE. Note that animal experiments of the DSP-CEST MRI were performed at a scanner with a small cavity in this study, the *B*_1_ field has limited inhomogeneities. However, the DSP-CEST signals need to be corrected by mapping the *B*_0_ inhomogeneity along with the *B*_0_ inhomogeneity when the proposed method is applied to the human studies, considering the *B*_1_ field could have substantial deviations in the ranges demonstrated in Fig. 4E-4H (57,58).

**Fig. 4.**
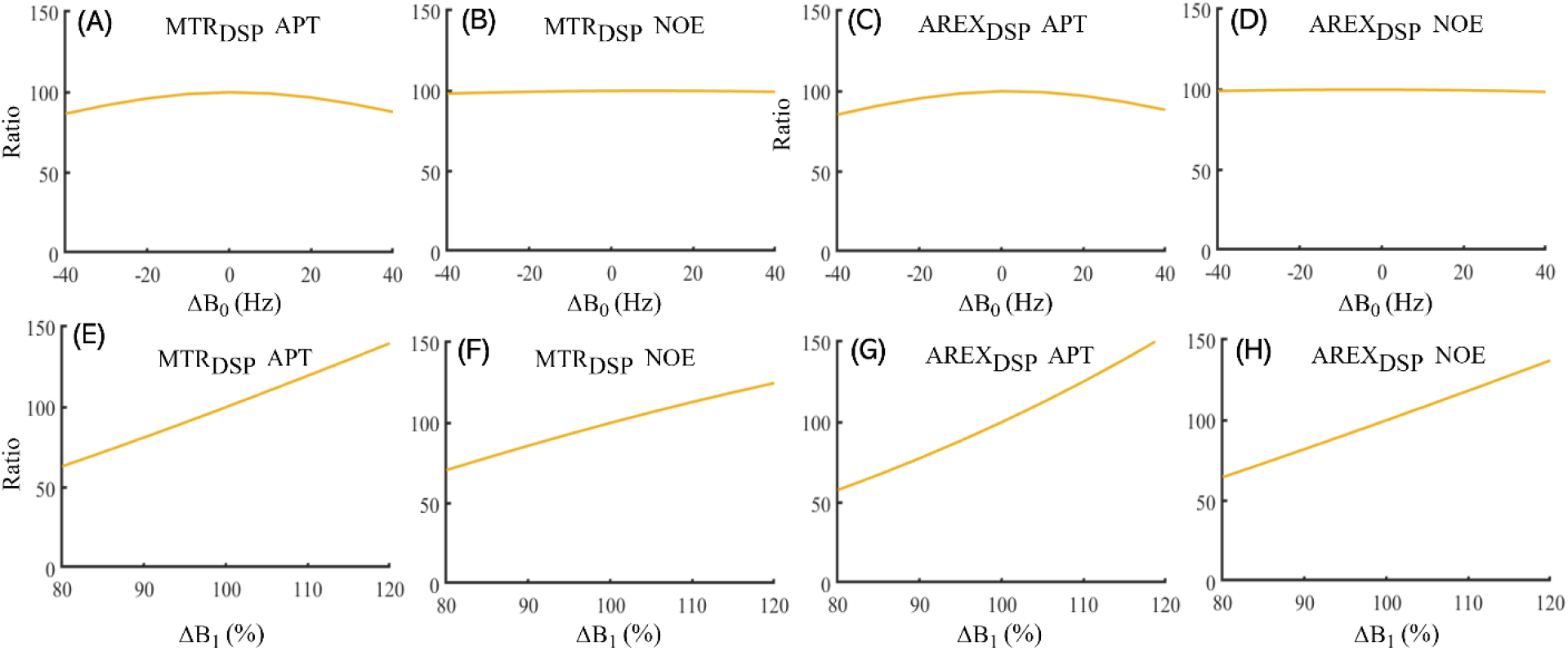
Influences of *B*_0_ inhomogeneity (Δ*B*_0_) and *B*_1_ inhomogeneity (Δ*B*_1_) on the DSP-CEST signals of the APT and the NOE. The simulations are performed with the *B*_0_ inhomogeneity spanning from −40 Hz to 40 Hz and the *B*_1_ inhomogeneity spanning from 80% to 120%, individually. Note that the *B*_1_ inhomogeneity is defined by the ratio of the deviated amplitude to the nominal value of RF field, and the influences of Δ*B*_0_ and Δ*B*_1_ are measured with the ratio of the value simulated with the inhomogeneity to the nominal value simulated without the inhomogeneity.

Fig. 5A-5D shows the conventional CEST Z-spectra with double saturation powers (B_1_L_ = 0.5 μT and B_1_H_ = 1 μT) and the corresponding DSP-CEST Z-spectra (including S_DPS_L_ and S_DPS_R_), which are obtained by averaging signals over the regions of tumors and contralateral normal tissues from three rats. Note that although the CEST signals (beyond ± 5ppm) in the raw CEST Z-spectra have considerable differences, the corresponding DSP-CEST signals match well in the Z-spectra, which is also observed in the simulations and suggests that the MT effects can be suppressed in the DSP-CEST metrics. Fig. 5E-5H shows the average MTR_DSP_ and AREX_DSP_ spectra from tumors and contralateral normal tissues of the three rats. The average MTR_LD_ and AREX_LD_ spectra were also plotted in Fig. 5E-5H to compare the LD-CEST method with the proposed DSP-CEST method. Note that the APT at 3.5 ppm, the guanidinium CEST at 2 ppm (an intermediate-exchange site), the NOE at −1.6 and −3.5 ppm can be observed in the spectra from both the LD analysis and the DSP-CEST. The MTR_DSP_ and AREX_DSP_ spectra are lower than the MTR_LD_ and AREX_LD_ spectra, which should be due to that the DSP method quantify a part of the CEST effect. The ratio of the signal at 2.0 ppm to the signal at 3.5 ppm in the DSP-CEST metrics is smaller than that in the LD-CEST metrics, which should be due to the exchange-rate low-pass filter effect of the DSP-CEST.

**Fig. 5.**
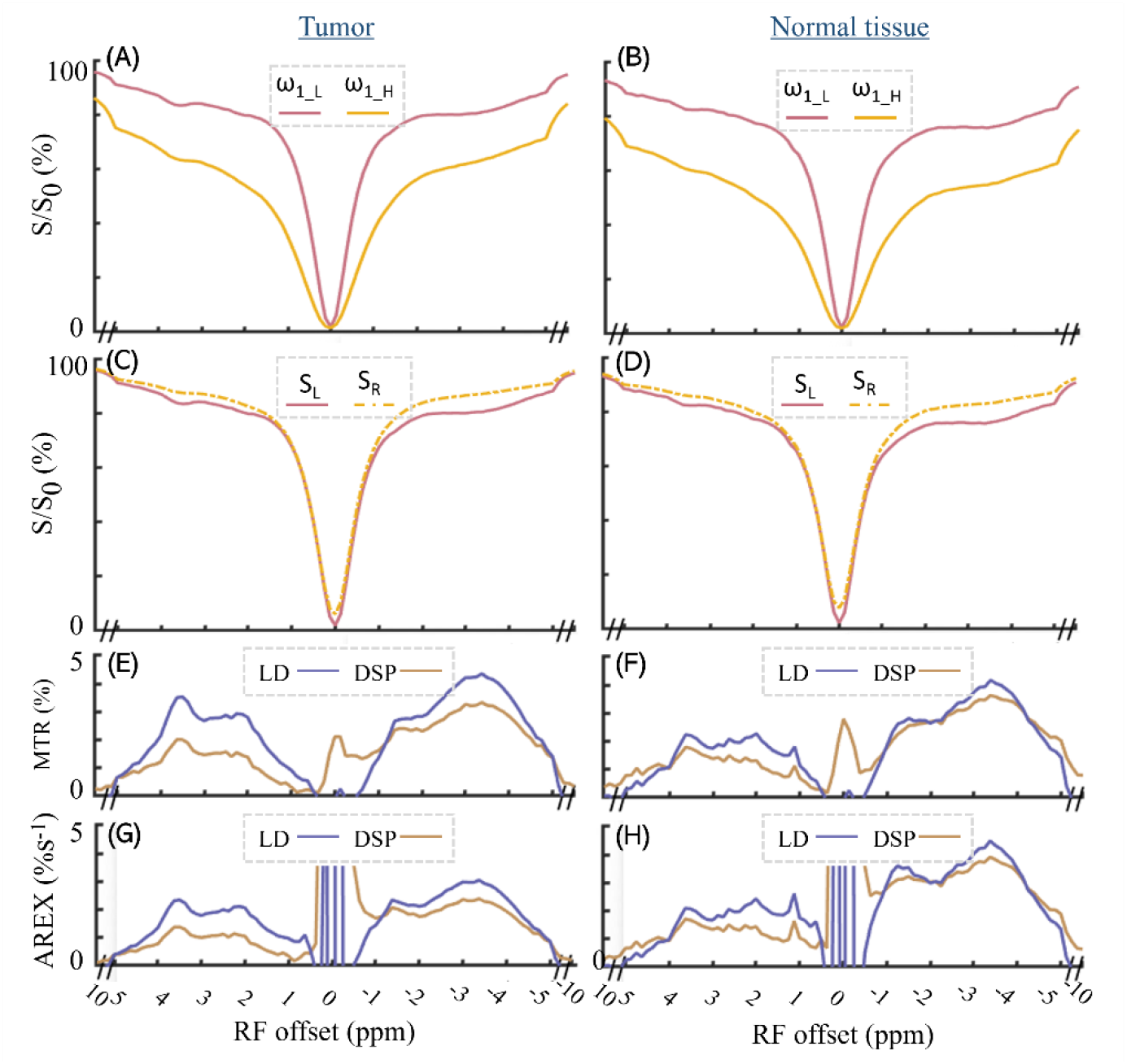
CEST Z-spectra with B_1_*L*_ = 0.5 μT and B_1_*H*_ = 1 μT (A, B), the corresponding S_DSP_L_ and S_DSP_R_ spectra (C, D), the MTR_DSP_ and MTR_LD_ spectra (E, F), and the AREX_DSP_ and AREX_LD_ spectra (G, H) obtained by averaging the corresponding signals over the regions of tumors (left column) and contralateral normal tissues (left column) from three rats.

Fig. 6 shows images of R_1w_, MT pool size ratio, APT and NOE(−3.5) that are acquired from a typical rat bearing tumors. The APT and NOE(−3.5) signals are determined with MTR_LD_, MTR_DSP_, AREX_LD_, and AREX_DSP_, respectively. The maps of R_1w_, MT pool size ratio, AREX_LD_ determined NOE(−3.5) and AREX_DSP_ determined NOE(−3.5) exhibit hypointense intensities in tumors while the map of MTR_LD_ determined APT shows hyperintense intensities in tumors. Furthermore, Fig. 7 shows results of statistical analysis based on ROIs that were delineated from the tumors and the contralateral normal tissues in the rat brains. Note that R_1w_, MT pool size ratio, MTR_LD_ determined APT, AREX_LD_ and AREX_DSP_ determined NOE(−3.5) have significant differences (*P* < 0.05) between the tumors and the normal tissues.

**Fig. 6.**
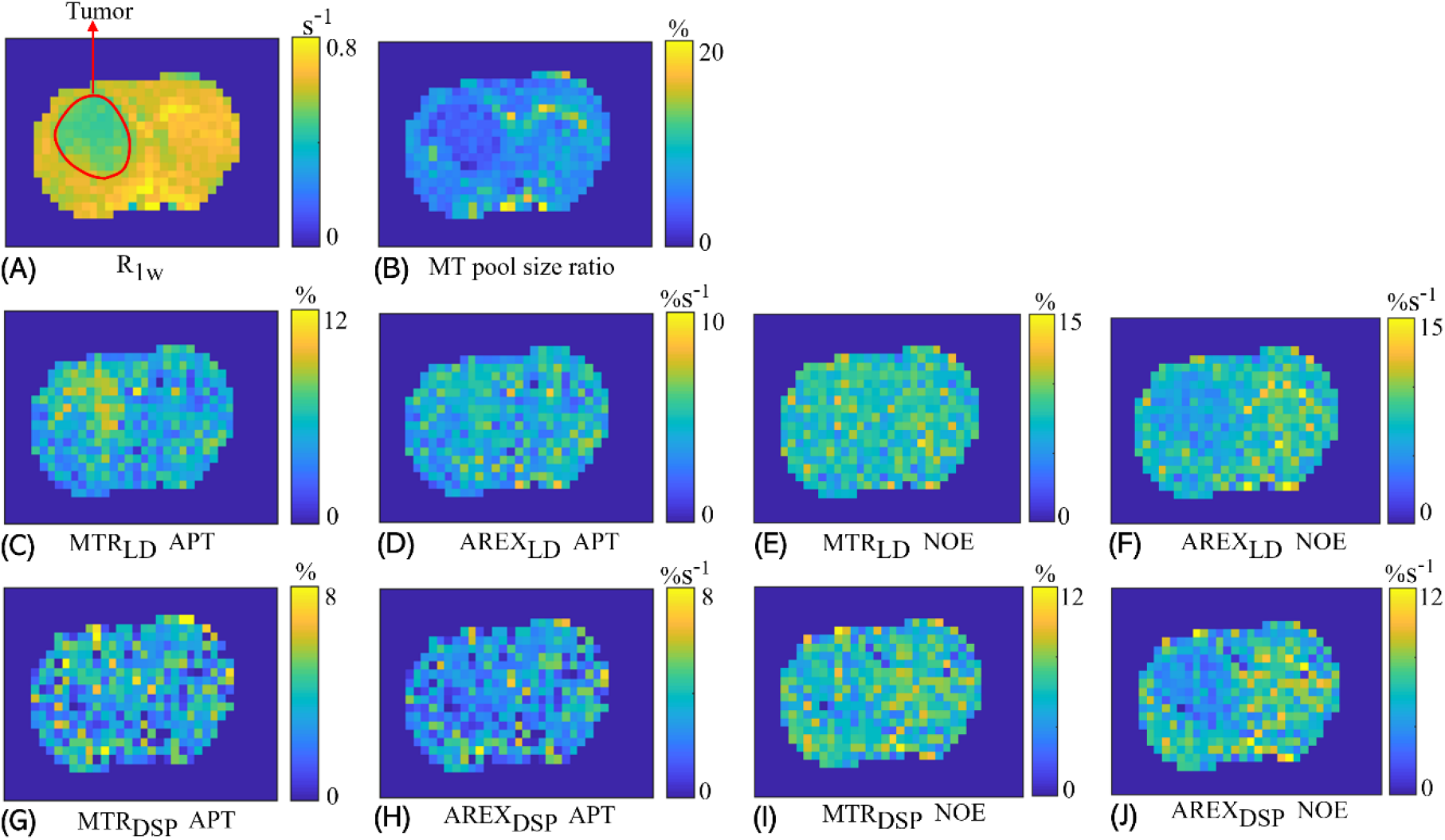
Images of R_1w_, MT pool size ratio and images of MTR_LD_, MTR_DSP_, AREX_LD_, and AREX_DSP_ of APT and NOE(−3.5) measured from a representative rat bearing tumors.

**Fig. 7.**
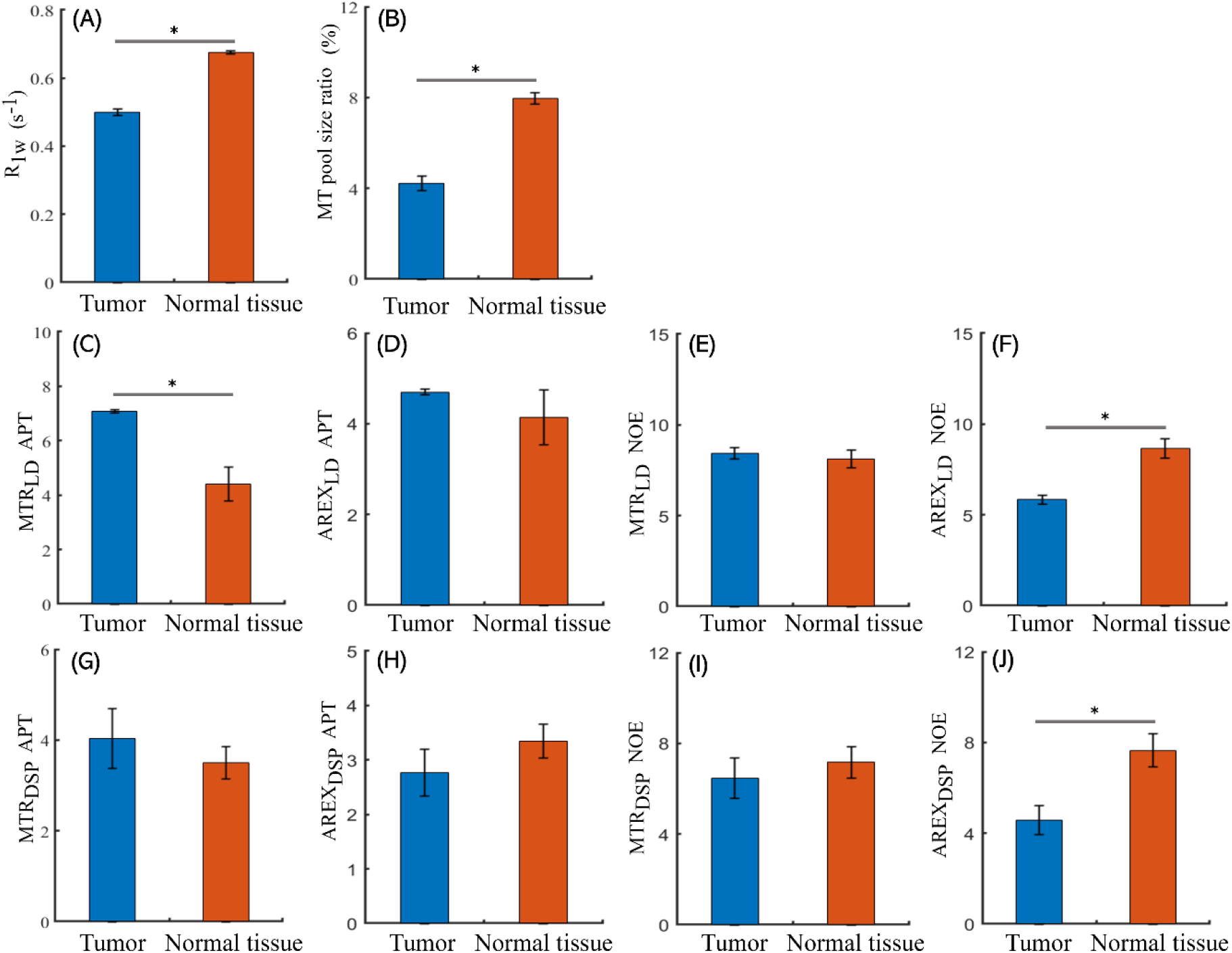
Comparisons between tumors and contralateral normal tissues for R1w, MT pool size ratio, and the APT and NOE(−3.5) signals quantified with MTR_LD_, MTR_DSP_, AREX_LD_ and AREX_DSP_. Note that the difference is deemed to be statistically significant if *P* < 0.05.

## DISCUSSION

In this study, a data-postprocessing method, termed DSP-CEST, was proposed to specifically quantify the APT and NOE effects based on the Z-spectra measured with conventional CEST sequences, where the signal acquired at a low saturation power is directly used for the label signal and the signal acquired at a high saturation power is processed by a well-tailored mathematical transformation to create a reference signal. By characterizing the differences between the label signal and the reference signal with two conventional metrics (MTR and AREX), the slow-exchange pool (i.e., amide and NOE(−3.5)) can be isolated from the background signals originating from the fast-exchange/coupling pool (i.e., lysine amine) and semi-solid MT, which is based on the different dependences of the pools on *ω*_1_. More specifically, at relatively low saturation powers, both the fast exchange CEST effect and the semi-solid MT effect increase linearly with 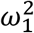, whereas the slow exchange CEST effect has no such relationship with 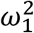. Thus, Eq. (9) is constructed to create a reference signal for the DSP-CEST, where the two undesired effects measured with different 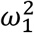 become almost equal after multiplying with a constant factor, and thus cancel with each other in the MTR and AREX metric. Although the data-postprocessing method is conducted with a CW-CEST sequence in this study, this method could also be available for the CEST sequences with the pulsed saturation RF if the pulsed RF is designed to remain the same relationship between the CEST effects of the three pools and the saturation power 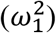.

The capability of the proposed method to specifically quantify the slow exchange/coupling effects is theoretically demonstrated with numerical simulations based on Bloch equations and further validated with *in vivo* experiments. As described above, the mathematical modeling could provide the interpretation of the proposed method with intuitive physical pictures using the mathematical derivation based on some approximations, but the deviations due to the approximations cannot be well estimated in the modeling and thus need to be further investigated using numerical simulations. In Fig. 2 and 3, signals from the fast exchange/coupling pool (i.e., lysine amine) and semi-solid MT are strongly suppressed in the simulated MTR and AREX, which demonstrate the validity of the approximations. In addition, the *in vivo* experiments show the two DSP-CEST Z-spectra (i.e., the label and reference signals) in Fig. 4c and 4d at chemical shifts beyond + 5 ppm (where there are only semi-solid MT effect) are very close, suggesting its ability to remove the semi-solid MT effects. (the small gap between the two DSP-CEST Z-spectra beyond −5ppm is due to the broad NOE(−3.5) effect). Furthermore, the ratio of the signal at 2.0 ppm (amide) to the signal at 3.5 ppm (guanidine in the intermediate exchange regime) in the MTR_DSP_ and AREX_DSP_ spectra is smaller than that in the MTR_LD_ and AREX_LD_ spectra, which should be due to the slow-exchange low-pass filter effect of the DSP-CEST, as shown in Fig. 3C, 3D, 3G and 3H. Note that, some substantial signals in the range of chemical shifts from 2 ppm to 3 ppm, which may originate from other slow-exchange pools besides the APT, such as phosphocreatine at 2.5ppm (55).

The *in vivo* experiments also demonstrate the ability of the proposed method to image the tumors in rats. Results of the experiments show that the NOE(−3.5) signals determined with the AREX_LD_ and AREX_DSP_ are significant different between tumors and contralateral normal tissues and the APT signals determined with the two metrics are not, which agrees with the results of previous reports observed with other methods (59,60). Besides, significant differences between tumors and contralateral normal tissues are detected by the MTR_LD_ determined APT, and not by the MTR_DSP_ determined APT. This may be due to the different exchange-rate selectivity of the two methods (i.e., the MTR_LD_ determined APT signal may detect signals from some fast-exchange pools).

Compared with conventional CEST quantification methods, the pros and cons of the proposed DSP-CEST methods are analyzed as below. (1) compared with conventional data fitting methods, the DSP-CEST does not requires assumptions of the number of the pools in the mode and the chemical shift and the line shape of each pool, and thus should be more robust. The multiple CEST/NOE peaks in the DSP-CEST in Fig. 5C-5H demonstrates its advantage over the multiple-pool Lorentzian fit. The LD analysis can also extract the multiple peaks. But it still needs to model the DS and semi-solid MT effect using a Lorentzian line function. However, the semi-solid MT in biological tissues is not a Lorentzian function but a super-Lorentzian function (57). In addition, the NOE(−3.5) may extend to beyond −5ppm and the fast exchange amine may extend to beyond +5ppm. CEST signals beyond these frequency offsets were used to fit the background DS and semi-solid MT effect which could causes incorrect fitting of the reference signal for the LD analysis. (2) the DSP-CEST method does not require the Z-spectrum acquisition within a broad coverage of chemical shifts that are necessary for the fitting method. A data acquisition within a narrow coverage of chemical shifts for a target site (APT or NOE) would substantially reduce the time cost of the acquisition. (3) the DSP-CEST does not rely on the development of a specific sequence and thus can be more favorable for clinical scanners. Although the CW-CEST sequence (that is not suitable for some clinical scanners) is used in this study, as analyzed above, a pulsed-CEST sequence that can be adapted for the DSP-CEST method by using pulsed-CEST signals acquired with two average saturation powers as we defined previously (61). (4) the DSP-CEST has a low signals contrast since it detects a part of the APT and NOE effect, which is shown in Fig. 3 and 5. The resulted decrease of SNR can be compensated by increasing average numbers of image acquisition. (5) the DSP-CEST is sensitive to the inhomogeneity of *B*_1_ (Fig. 4). Hence, the DSP-CEST signals need to be corrected by mapping the *B*_1_ inhomogeneity when the DSP-CEST method is applied to the human studies, where the *B*_1_ field could have substantial deviations.

## CONCLUSION

In this study, we proposed a new data-postprocessing method to quantify the APT and NOE effects with considerably increased specificities and a reduced cost of imaging time.

## APPENDIX A

A two-pool (water pool and semi-solid MT pool) coupled Bloch equation has been used to describe the MT effect (62). The solution for this two-pool model has been derived to be (63),

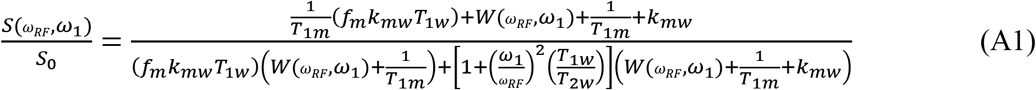

where *k_mw_* and *k_mw_* are the MT rate between water pool and the semi-solid MT pool longitudinal relaxation time, respectively. 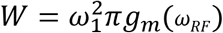, which is the saturation rate of the semi-solid MT pool. *g_m_* is a super-Lorentzian absorption line shape in biological tissues (63–65).

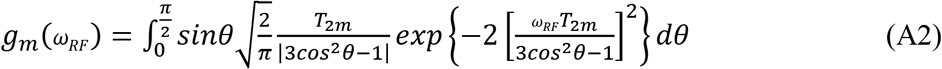

where *T*_2*m*_ is the semi-solid MT pool transverse relaxation time.

Eq. (A1) can be rewritten as,

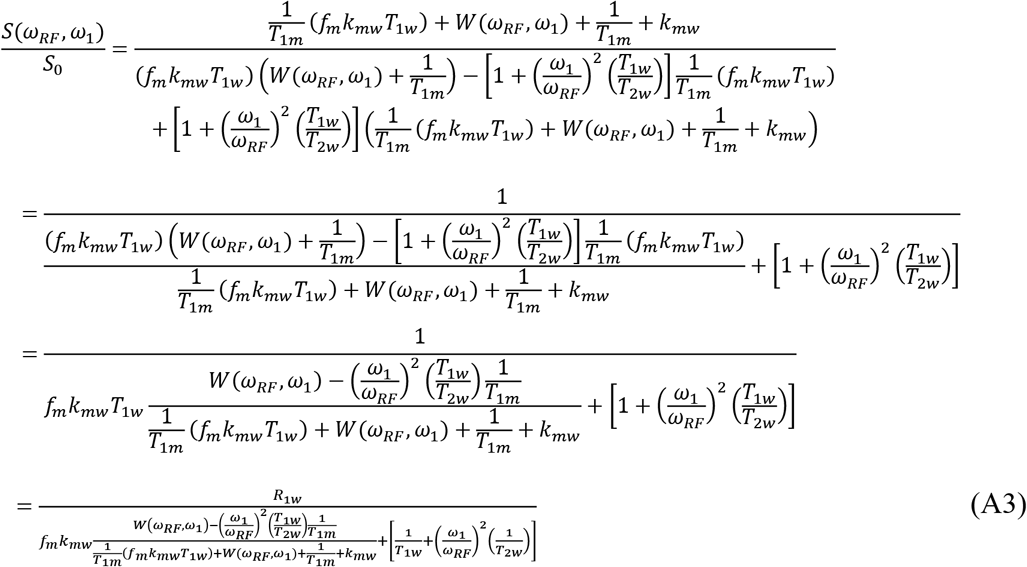

Since *ω_RF_* is much higher than *ω*_1_, 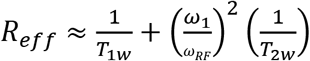. By comparison with 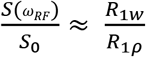 derived from Eq. (2), the first term in the denominator of Eq. (A3) should be 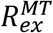 in a two-pool (water and semi-solid MT pool) model.

Since *k_mw_* is much higher than 1/*T*_1*m*_ and (*f_m_k_mw_T*_1*w*_)/ *T*_1*m*_, 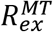 can be simplified to,

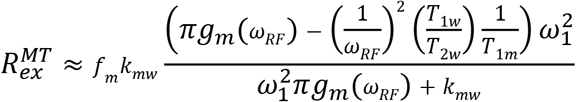

When 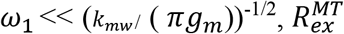 is proportional to 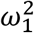. For *k_mw_* = 25 s^−1^ (66), *T*_2*m*_= 15 μs (66), *ω_RF_* = ±3.5 ppm at 4.7 T, and a symmetric super-Lorentzian absorption line shape, (*k_mw_*/(*πg_m_*))^−1/2^/3≈1 μT.

